# Novel and known transcriptional targets of ALS/FTD protein TDP-43: Meta-analysis and interactive graphical databases

**DOI:** 10.1101/2021.12.08.471595

**Authors:** Maize C. Cao, Emma L. Scotter

## Abstract

TDP-43 proteinopathy is the major pathological hallmark of amyotrophic lateral sclerosis (ALS) and tau-negative frontotemporal dementia (FTD). Mounting evidence implicates a loss of normal TDP-43 function in neurodegeneration, either resultant from or independent of TDP-43 aggregation. TDP-43 knockdown is therefore a common paradigm for modelling ALS and FTD. However, because TDP-43 can interact directly with thousands of mRNA targets and regulate the function of other RNA binding proteins, the phenotype of TDP-43 depletion is likely to differ depending on the proteomic and transcriptomic profile of the model cell type. Here, we conducted a meta-analysis of publicly available RNA-sequencing datasets that utilized TDP-43 knockdown to model ALS or FTD, and validated these against RNA-sequencing data from TDP-43-immunonegative neuronal nuclei from ALS/FTD brain. We present these analyses as easy-to-use interactive graphical databases. Of 9 TDP-43-knockdown datasets identified, 4 showed significant depletion of TARDBP (human HeLa and SH-SY5Y cell lines, induced human motor neurons, and mouse striatal tissue). There was little overlap in differentially expressed genes between TDP-43-knockdown model cell types, but *PFKP, RANBP1, KIAA1324, ELAVL3*, and *STMN2* were among the common TDP-43 targets. Similarly, there were few genes that showed common patterns of differential exon usage between cell types and which validated in TDP-43-immunonegative neurons, but these included well-known targets *POLDIP3, RANBP1, STMN2*, and *UNC13A*, and novel targets *EXD3, CEP290, KPNA4*, and *MMAB*. Enrichment analysis showed that TDP-43 knockdown in different cell types affected a unique range of biological pathways. Together, these data identify novel TDP-43 targets, validate known TDP-43 targets, and show that TDP-43 plays both conserved and cell-type-specific roles in the regulation of gene expression and splicing. Identification of cell-type-specific TDP-43 targets will enable sensitive mapping of cell-autonomous TDP-43 dysfunction beyond just neurons, while shared TDP-43 targets are likely to have therapeutic value across myriad cell types.

## Introduction

TDP-43 is a nuclear DNA- and RNA-binding protein first discovered to bind to the trans-active response element in the HIV-1 sequence (1). TDP-43 was subsequently found to be the major constituent of pathogenic aggregates in amyotrophic lateral sclerosis (ALS) and frontotemporal dementia (FTD) neuropathology (2, 3). Indeed, hyper-phosphorylated and aggregated cytoplasmic TDP-43 is the pathological signature in almost all cases of ALS, and approximately 50% of FTD patients (3-5). Other neurodegenerative diseases also manifest with TDP-43 neuropathology, including Alzheimer’s disease, Parkinson’s disease and Huntington’s disease (6-13). There is also a clear relationship between the development of neurodegenerative disease and mutations in other RNA-binding proteins that favor their aggregation (14-17). Strikingly, the regional patterning of neuronal loss both in the brain and spinal cord closely reflects the patterning of TDP-43 aggregate deposition (18-20). However, TDP-43 protein inclusions represent only one species across a spectrum of conformations in which TDP-43 can exist (21) and their ease of detection has likely influenced our perception of the pathomechanisms of disease.

The gain-of-toxic-function hypothesis for TDP-43 in ALS emerged from three key findings; i) that almost all inherited ALS including *TARDBP*-mutant ALS is inherited dominantly (22, 23); ii) that the characteristic pathology is the appearance of cytoplasmic TDP-43 aggregates absent from non-neurodegenerative disease tissue (2, 3); and iii) that TDP-43 overexpression paradigms in animal models recapitulated these TDP-43 aggregates and the symptoms of human ALS (24-27). Recognition that a loss of normal TDP-43 function may also, or instead, be pathogenic in ALS was based upon the observation that TDP-43, both in human ALS tissue and transgenic animal models harboring TDP-43 inclusions, is cleared from its normal location in the nucleus (28-31). Nuclear-to-cytoplasmic mislocalization of TDP-43 likely feeds into, and is further induced by, TDP-43 aggregation in the cytoplasm via a sequestration mechanism (30, 32-34) (Fig. 1). Although TDP-43 fulfils specific cytoplasmic functions, including the regulation of stress granules which are proposed by some to ‘seed’ TDP-43 aggregation (35-38), TDP-43 is a predominantly nuclear protein and the majority of its RNA processing roles are executed in the nucleus (1, 39-47). These include regulation of alternative splicing, enhancement or repression of exon and cryptic exon inclusion, mRNA transport, and polyadenylation (46-53). Nuclear TDP-43 is also able to autoregulate its own mRNA levels through a negative feedback loop by binding its own 3’ UTR (54), so loss of TDP-43 from the nucleus likely further contributes to TDP-43 overproduction, phase separation and aggregation, and sequestration (55) (Fig. 1). Loss of appropriate TDP-43 RNA processing function is evidenced in human ALS by extensive transcriptional change and mis-splicing (47, 48, 53, 56). Notably, similar transcriptional changes, motor neuron pathology, and motor symptoms are seen both in animal models with TDP-43 inclusions, and in models without TDP-43 inclusions which are based upon TDP-43 knockdown (57, 58). Thus, loss of nuclear TDP-43 function is clearly critical to the pathogenesis of ALS.

**Figure 1.**
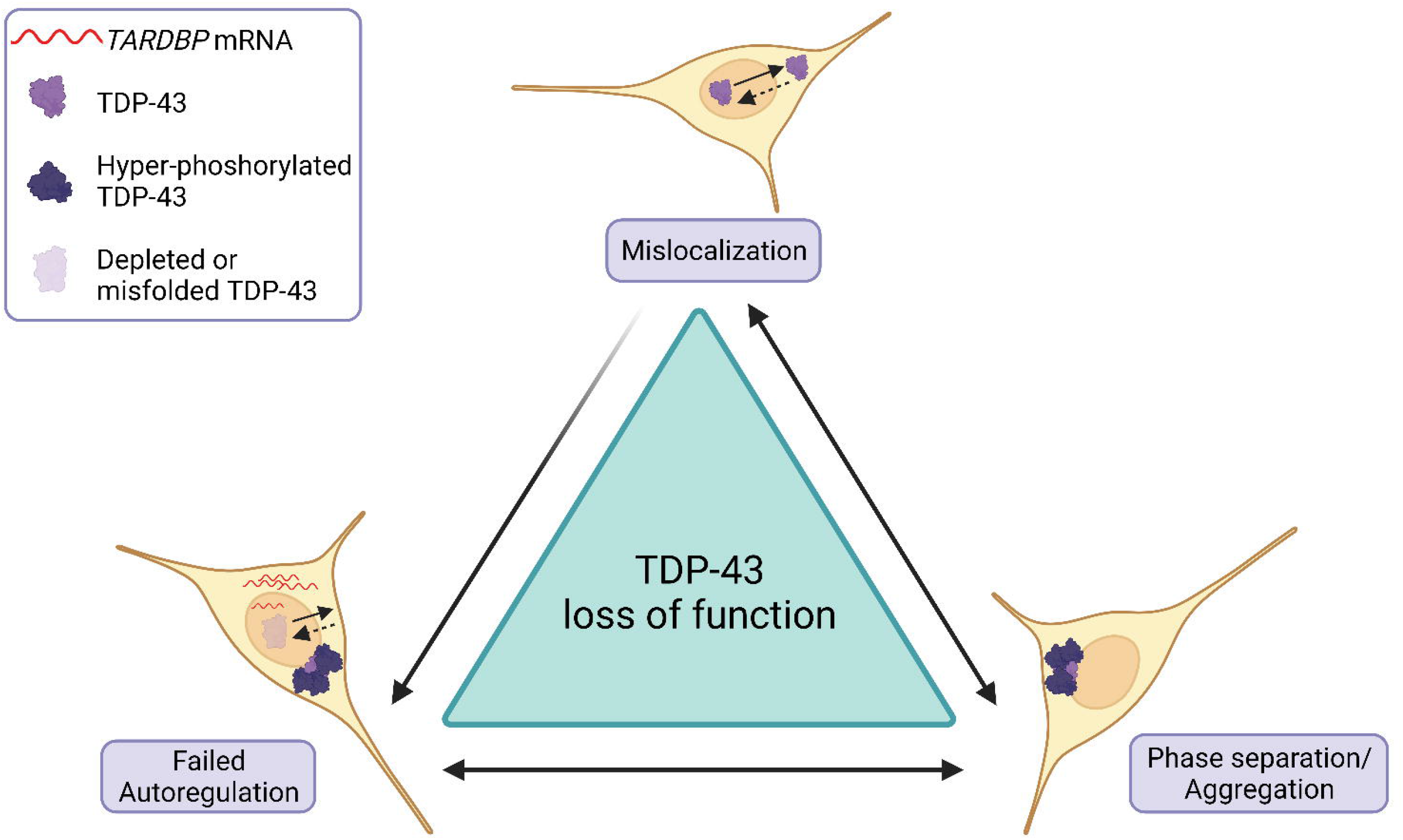
Schematic of the mechanisms of TDP-43 loss of function in ALS. *Top*. TDP-43 is actively imported into the nucleus and passively diffuses (132), giving TDP-43 its predominantly nuclear localization. However, TDP-43 in ALS is frequently mislocalized to the cytoplasm, leading to nuclear TDP-43 depletion. *Bottom right*. Cytoplasmic TDP-43 is prone to phase separation and aggregation, with aggregates becoming hyperphosphorylated and further sequestering TDP-43. *Bottom left*. Readily detectable TDP-43 nuclear depletion or sequestration into aggregates, or less easily detected misfolding, can deplete the functional pool of TDP-43. TDP-43 loss of function leads to failed autoregulation of *TARDBP* (enhancing TDP-43 translation), in addition to failed regulation of myriad other TDP-43 targets many of which are unknown or unvalidated.

Recognizing the impact that TDP-43 loss-of-function may have on gene expression in ALS and FTD, an increasing number of studies report the transcriptome-wide effect of TDP-43 depletion. While certain TDP-43 targets such as *RANBP1* and *POLDIP3* are clearly reproducible in multiple studies (43, 44, 46, 53, 57, 59, 60), there has yet to be a formal analysis published of common TDP-43 targets. Different cell types are obviously transcriptomically unique, as are the same cell types derived from different species, meaning the influence of TDP-43 on gene expression and splicing is context-dependent (46, 61). Here, we re-analyzed publicly available RNA-sequencing datasets from TDP-43-depleted model systems to examine common and unique transcriptional patterns of TDP-43 loss. We then examined whether TDP-43 targets common to these experimental TDP-43 depletion datasets were recapitulated in a human ALS brain dataset. Elucidating markers of TDP-43 loss-of-function will enable better understanding of disease mechanisms and the extent to which TDP-43 loss-of-function is associated with neurodegeneration. Further, such markers may serve as biomarkers and/or targets for treatment.

## Methods

### Identification of TDP-43 knockdown studies for meta-analysis

A repository search was conducted in September 2020 to identify TDP-43 knockdown studies with available RNA-seq data using the Gene Expression Omnibus (GEO) from the National Centre for Biotechnology Information (NCBI, http://ncbi.nlm.nih.gov/geo). The search was performed using the keyword “TDP-43” and the results were filtered by setting Entry Type as ‘Series’ to capture all potential samples that belonged to a common study and setting Study Type as ‘Expression profiling by high throughput sequencing’. Inclusion criteria for re-analyzing these datasets were: 1) Raw RNA-seq data available; 2) At least three TDP-43 knockdown samples and two appropriate control samples; 3) Experimental depletion of TDP-43. In addition, an RNA-seq dataset was identified in which neuronal nuclei had been sorted from ALS/FTD tissue according to nuclear TDP-43 immunoreactivity; either TDP-43-positive (TDPpos; normal nuclear TDP-43) or TDP-43-negative (TDPneg; mislocalized TDP-43) (62). This was considered an appropriate ‘disease validation’ dataset for targets identified through meta-analysis of TDP-43 knockdown studies, because the within-case comparison of TDP-43-positive and -negative neuronal nuclei paralleled the paradigm of TDP-43 loss of function by knockdown (Table 1, Fig. 2).

**Figure 2.**
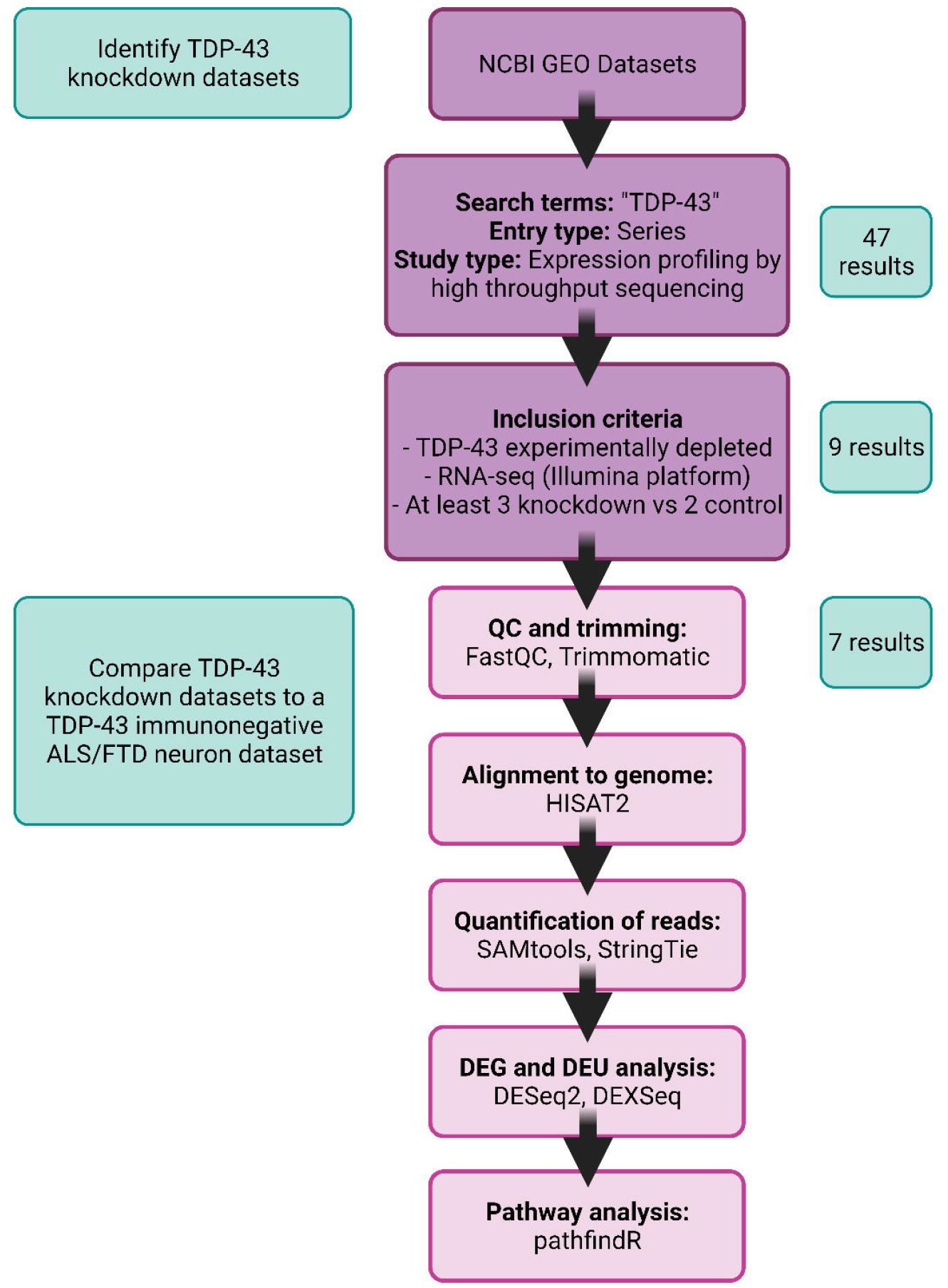
Study selection and RNA-seq data processing pipeline. RNA-seq datasets were selected from the NCBI GEO database using search term and study type, filtered for studies that performed RNA-Seq on models with experimental TDP-43 depletion, and processed through a common bioinformatic pipeline (FastQC, Trimmomatic, HISAT2, SAMtools, StringTie, DESeq2, DEXSeq and pathfindR). DEG: differentially expressed genes; DEU: differential exon usage. TDP-43 knockdown datasets were then compared with a TDP-43-immunonegative ALS/FTD neuronal nuclei dataset.

### Data processing and statistical analysis

Raw data were downloaded from NCBI using Sequence Read Archive (SRA) Toolkit (http://www.ncbi.nlm.nih.gov/sra), and quality control was applied using FastQC software (63). Samples were assessed based on per base sequence quality, with the threshold set at Phred score of at least 30. Samples were excluded from analysis if they had no reads that survived trimming. If this reduced the sample number of the study to below that of the inclusion criterion, the study was excluded. Raw data that still contained adapter sequence content were trimmed with Trimmomatic 0.39 (64). Code for trimming is included in supplementary file S1. Reads were then aligned to the appropriate reference genome with HISAT2 (65) and quantified with StringTie (66). Reference genome builds used were GRCh38, GRCm39, and Rnor6.0 for human, mouse, and rat respectively. Count data was imported using R package tximport (67) to perform differential expression analysis using DESeq2 (68) and differential exon usage analysis using DEXSeq (69). Differential exon usage is a more general measure than alternative splicing, as differing exon boundaries between reads are accounted for, therefore revealing differential usage of parts of exons or introns (exonic elements) (69). *DESeq2* p-values were calculated using Wald tests and corrected for multiple testing using the Benjamini-Hochberg method. Outliers were determined by Cook’s distance. Genes with adjusted p-values <0.05 were considered significantly changed and differential exon usage was considered significant with adjusted p-value <0.1. R package *pathfindR* (70) was used for enrichment analysis using the Reactome gene set (https://reactome.org/). Pathways with p-values <0.05 were used for ‘parent’ pathway visualization.

Interactive graphical databases generated from the packages *‘Glimma’* and *‘DEXSeq’* can be accessed via GitHub (Supplementary file S2). These enable the reader to visualize and interact with all differential gene expression and exon usage analyses described herein, including genes of interest not highlighted in our study.

### Data visualization

Data was visualized using R software with *‘ggplot2’* (71), *‘ggvenn’* (72), *‘DESeq2’* (68), and *‘DEXSeq’* (73), Prism 9.0 software (La Jolla, CA), and Integrative Genomics Viewer (IGV) (74). Adobe Photoshop 2021 22.5.1 (Adobe Inc.) was used as a graphic editor.

## Results

### Validation of TARDBP depletion in TDP-43 knockdown models and in TDPneg ALS/FTD neuronal nuclei

Forty-seven RNA-sequencing datasets were identified using “TDP-43” as a keyword. Nine studies met the inclusion criteria for re-analysis (raw data available, appropriate sample size, TDP-43 depleted experimentally (“knockdown”)), however only seven of these met quality control thresholds and were fully processed (Fig. 2). Of these seven, three were performed on cells derived from humans, three from mouse, and one from rat (Table 1). An additional RNA-sequencing dataset was identified in which neuronal nuclei had been sorted from ALS/FTD tissue according to nuclear TDP-43 immunoreactivity. This dataset also met quality control thresholds and was processed using the same pipeline (Fig. 2).

Following processing of raw data, *TARDBP* mRNA levels were plotted with DESeq2 *‘plotCounts’* function to confirm TDP-43 knockdown at the RNA level (Fig. 3). Significant *TARDBP* knockdown was seen in studies GSE27218 (mouse striatum) (53), GSE136366 (human HeLa cells) (59), GSE122069 (human SH-SY5Y cells) (52), and GSE121569 (human induced motor neurons, ihMN) (51) (Fig. 3A). In contrast, TARDBP levels were not significantly different between control and knockdown samples for studies GSE21993 (mouse embryonic stem cells) (75), GSE116456 (mouse mammary gland) (76), and GSE135611 (rat primary astrocytes) (77) by this methodology (Fig. 3B), so these studies were not further analyzed. Finally, in study GSE126543 (cortical neuronal nuclei with or without detectable TDP-43 immunolabelling from seven ALS/FTD human brains) (62), two samples showed striking loss of *TARDBP* from TDP-43-immunonegative nuclei (s2, s7), three showed subtle loss of *TARDBP* (s1, s5, s6), and two showed increased *TARDBP* (s3, s4; Fig. 3C).

**Figure 3.**
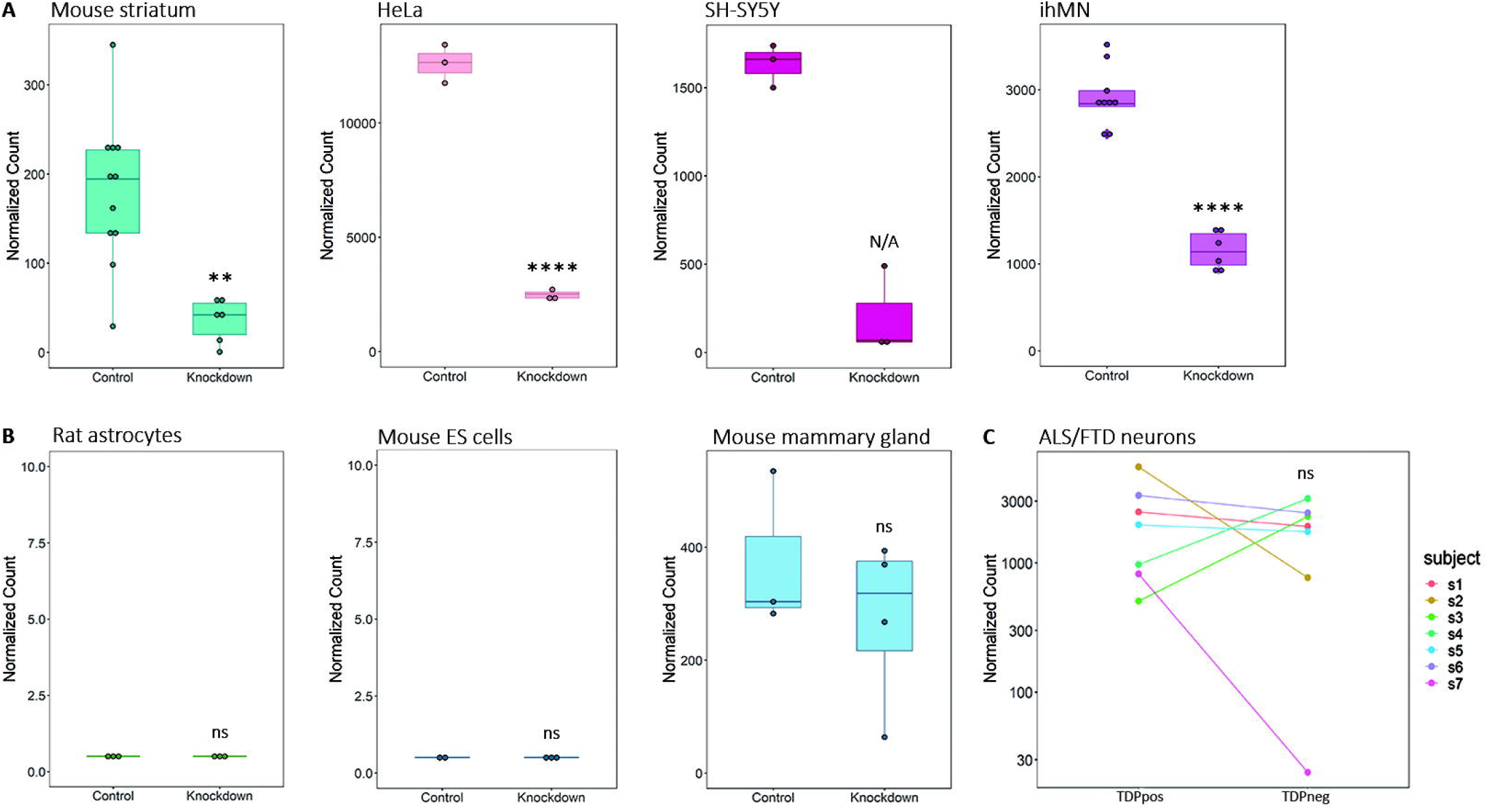
TARDBP depletion in TDP-43 knockdown models and in TDPneg ALS/FTD neuronal nuclei. A-B. Normalized counts of *TARDBP* transcript from TDP-43 knockdown studies. A. Studies included in the meta-analysis, from left to right - GSE27218 (mouse striatum), GSE136366 (HeLa), GSE122069 (SH-SY5Y), GSE121569 (ihMN). *Bottom row*. B. TDP-43 knockdown studies not included in the meta-analysis, from left to right - GSE135611 (rat astrocytes), GSE21993 (mouse embryonic stem cells), GSE116456 (mouse mammary gland). C. Normalized counts of subject-paired *TARDBP* transcript in TDP-43-immunopositive (TDPpos) ALS/FTD and TDP-43-immunonegative (TDPneg) ALS/FTD neuronal nuclei (GSE126542). Ns, not significant; **, padj <0.005; ****, padj <0.00005; N/A, DESeq2 detected an extreme outlier count by Cook’s distance.

### Shared, cell-type-specific, and species-specific differentially expressed genes with TDP-43 knockdown and in TDPneg ALS/FTD neuronal nuclei

Of the four knockdown studies with adequate *TARDBP* depletion, three models were derived from human (HeLa, SH-SY5Y, ihMN) and one from mouse (striatum). Of these, ihMN, SH-SY5Y, and mouse striatum comprised or included cells with a neuronal phenotype. DESeq2 analysis identified sets of differentially expressed genes following *TARDBP* knockdown for each dataset (Supplementary file S3), interactive visualization of which is presented using the *Glimma* package (Supplementary file S2). In contrast, DESeq2 analysis in the TDP-43 negative versus positive nuclei dataset did not indicate *TARDBP* to be significantly decreased, but selected genes of interest were examined as described below.

Differentially expressed gene sets were compared between TDP-43 knockdown datasets, revealing no significantly changed genes (padj <0.05) shared by all four studies, five genes shared by three studies (*TARDBP, PFKP, MAFB, FKBP4, AL109811*.*3*), and 46 genes shared by two studies (Fig. 4A, Supplementary file S3). Nine genes of interest, as indicated in Fig. 4A, were selected for further examination.

**Figure 4.**
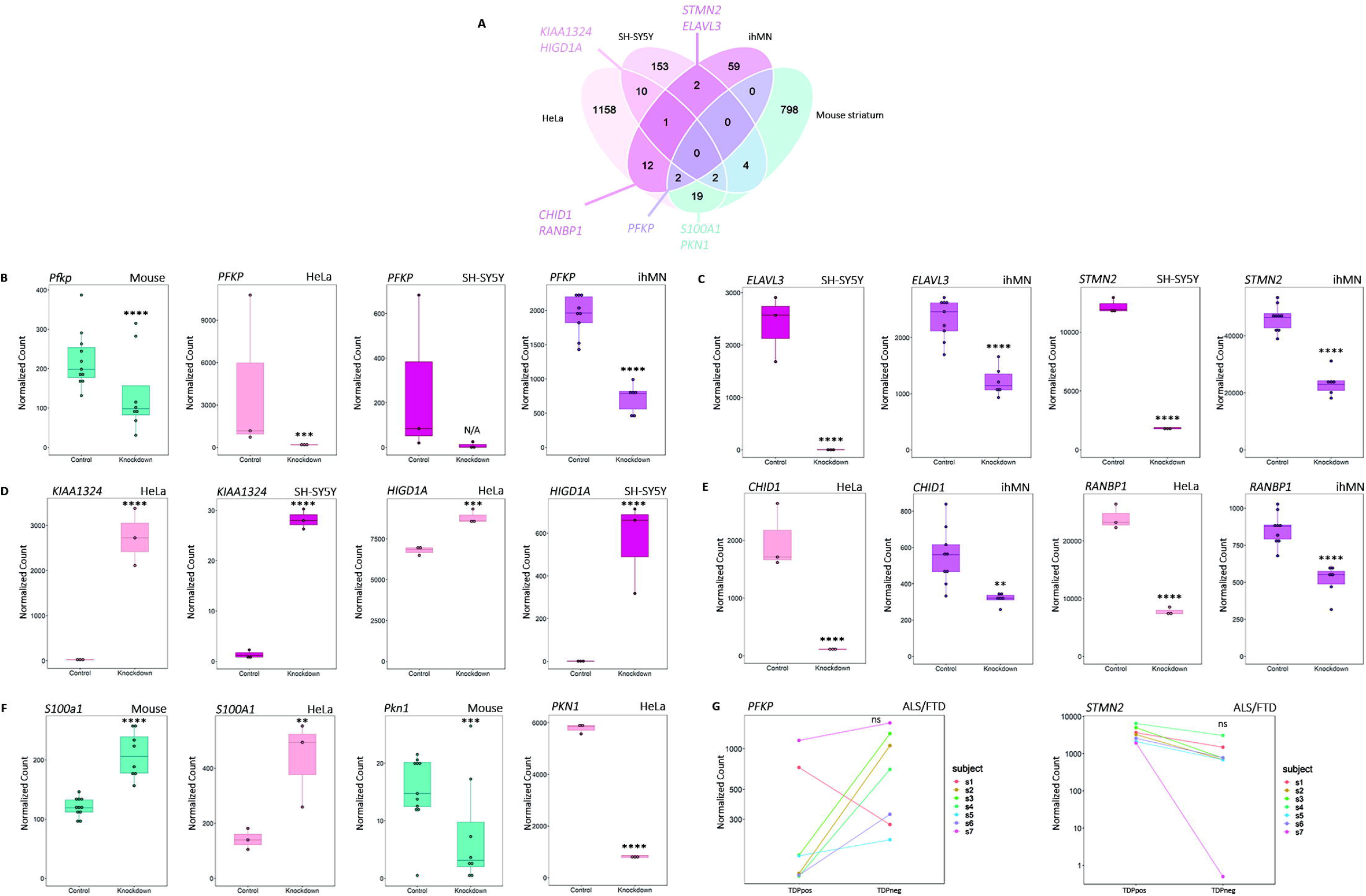
Shared, cell-type-specific, and species-specific differentially expressed genes with TDP-43 knockdown and in TDPneg ALS/FTD neuronal nuclei. A. Venn diagram of differentially expressed genes (DEGs) common between HeLa, SH-SY5Y, ihMN and mouse striatum. B. Normalized counts of *PFKP* transcript from *mouse striatum*, HeLa and SH-SY5Y samples. C. Normalized counts of *ELAVL3* and *STMN2* transcript from SH-SY5Y and ihMN samples. D. Normalized counts of *KIAA1324* and *HIGD1A* transcript from HeLa and SH-SY5Y samples. E. Normalized counts of CHID1 and *RANBP1* transcript from HeLa and ihMN samples. F. Normalized counts of *S100A1* and *PKN1* transcript from mouse striatum and HeLa samples. G. Normalized counts of *PFKP* and *STMN2* transcript from ALS/FTD TDP-43-immunopositive (TDPpos) and TDP-43-immunonegative (TDPneg) ALS/FTD neuronal nuclei. Ns, not significant; **, padj <0.005; ***, padj <0.0005; ****, padj <0.00005; N/A, DESeq2 detected an extreme outlier count by Cook’s distance.

*PFKP* (phosphofructokinase, platelet type) was the most conserved TDP-43 target, being significantly decreased in three knockdown studies and observably lower in the fourth study, and with the log2FC for each study being less than -1 (greater than 2-fold decrease) (Fig. 4B). A decrease in *PFKP* was only seen in TDPneg ALS/FTD neuronal nuclei for one sample (s1), with all other samples showing *PFKP* to increase (Fig. 4G). Two notable DEGs that were decreased in cell types with human neuronal phenotype (SH-SY5Y and ihMN) were *STMN2* and *ELAVL3* (Fig. 4C). *STMN2* encodes stathmin-2 which regulates microtubule dynamics in neurons (78), and *ELAVL3* encodes ELAV-like protein 3 which is a neuron-specific RNA-binding protein (79). Both have been reported to be TDP-43 targets (51, 52). A decrease in *STMN2* (Fig. 4G) but not *ELAVL3* (not shown), was also observed in TDPneg ALS/FTD neuronal nuclei in all samples, with near complete suppression of *STMN2* expression in s7.

Other common DEGs between pairs of studies were selected based on fold changes of at least 1.5 in the same direction, and the most significant adjusted p-value from each sample. These were *KIAA1324* and *HIGD1A* (between HeLa and SH-SY5Y, Fig. 4D), *CHID1* and *RANBP1* (between HeLa and ihMN, Fig. 4E) and *S100A1* and *PKN1* (between HeLa and mouse striatum, Fig. 4F). However, none of these was significantly changed in TDPneg ALS/FTD neuronal nuclei (not shown).

### Shared and unique biological pathways changed with TDP-43 knockdown

Having identified genes with conserved patterns of regulation by TDP-43 between cell types and species, we examined the effect of TDP-43 knockdown on biological pathways. TDPneg ALS/FTD neuronal nuclei were not included in pathway analysis because of the lack of significant DEGs as described above. Each TDP-43 knockdown cell type displayed a different makeup of changed pathways (Fig. 5, Supplementary file S4). For HeLa cells, the most changed parent pathways were gene expression (transcription) and extracellular matrix organization (Fig. 5A). For SH-SY5Y, the most changed parent pathways were signal transduction and cellular response to external stimuli (Fig. 5B); for ihMN, they were cell cycle and disease (Fig. 5C); and for mouse striatum, they were signal transduction and neuronal system (Fig. 5D). Signal transduction was therefore common to the three neuronal models, while cellular responses to external stimuli, disease, and gene expression (transcription) were common to human models. The heterogeneity of changed pathways in different cell types upon TDP-43 knockdown may be reflective of the diverse pathways implicated in disease.

**Figure 5.**
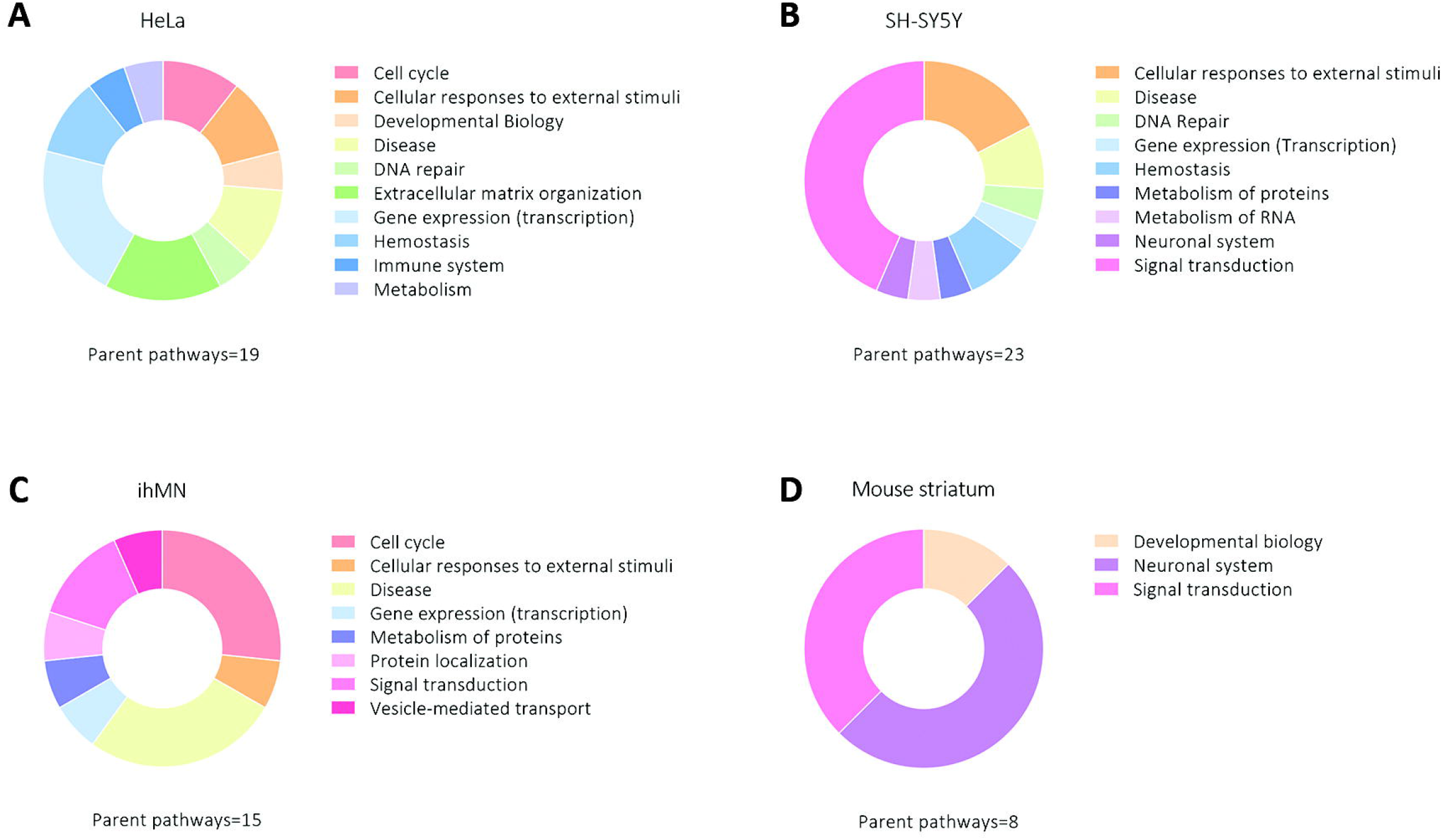
Shared and unique biological pathways changed with TDP-43 knockdown. Donut plots of the most changed biological pathway types with TDP-43 knockdown in A. HeLa, B. SH-SY5Y, C. ihMN, D. mouse striatum. Pathways significantly enriched in each cell type were assigned to parent pathways and grouped according to parent pathway type. The total number of parent pathways represented is given below each donut.

### Known differentially used exons with TDP-43 knockdown and in TDPneg ALS/FTD neuronal nuclei: shared, cell-type-specific, or species-specific

As TDP-43 is known to regulate splicing and cryptic exon suppression (46, 47), we next examined differential exon usage between control and TDP-43-knockdown samples for each study, and TDPpos and TDPneg ALS/FTD neuronal nuclei, using the *DEXSeq* package (Supplementary file S4). This package detects changes in the relative usage of exons or parts of exons between conditions and generates a graphical display of the differentially used exonic elements (Supplementary file S2). Genes with differentially expressed exonic elements were compared between TDP-43 knockdown datasets, revealing 2 genes shared by all four studies (*POLDIP3* and *TRAPPC12*), 34 genes shared by three studies (including *EXD3, KPNA4, MMAB, GOSR2*, and *HP1BP3*), and 253 genes shared by two studies (including RANBP1, CEP290, and STMN2) (Fig. 6A, Supplementary file S4). Ten genes of interest, as indicated in Fig. 6B, were selected for showcase for their consistent exonic changes. All except *TRAPPC12* showed the same differential exon/s between several TDP-43 knockdown samples and TDPneg ALS/FTD neuronal nuclei.

**Figure 6.**
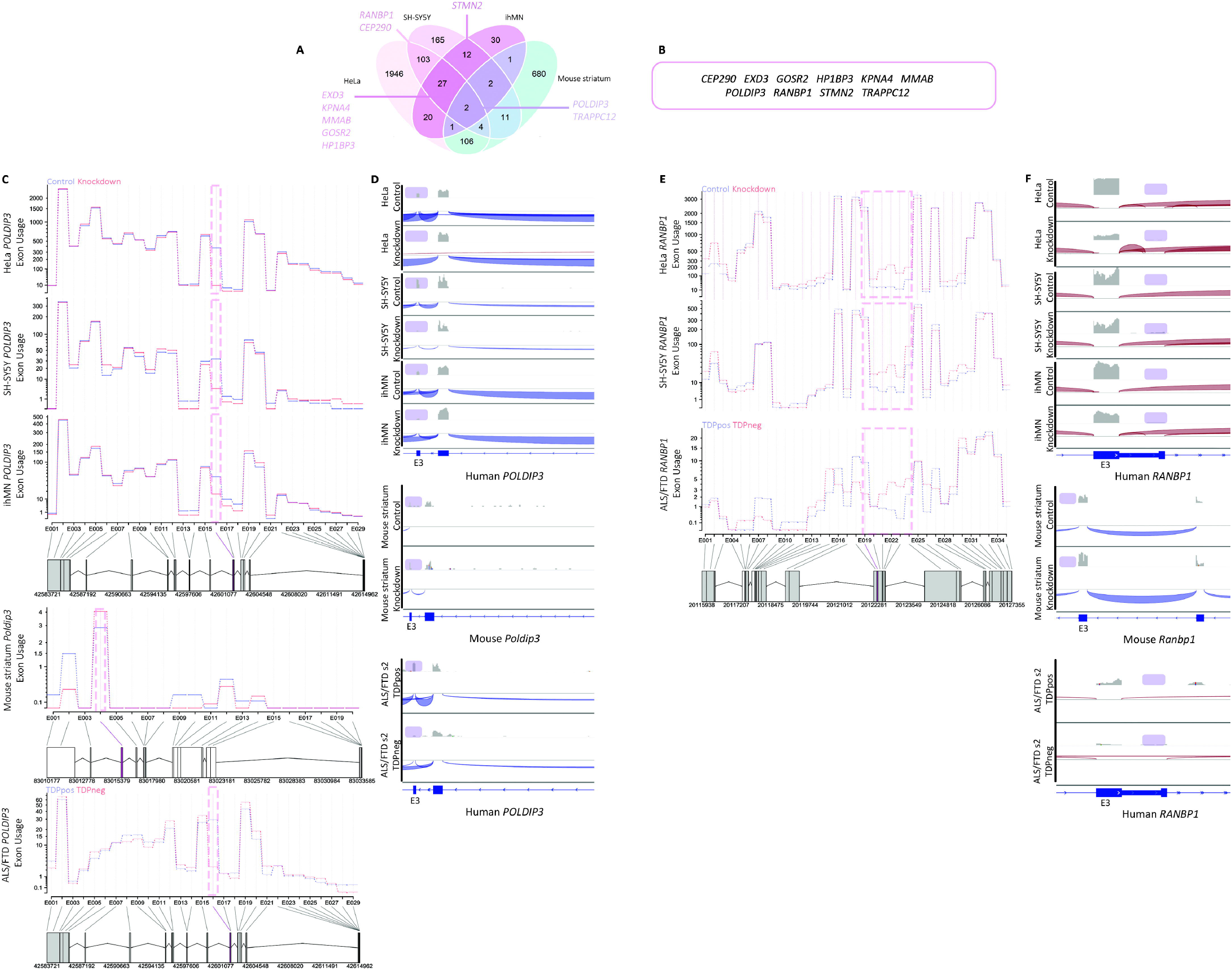
Known differentially used exons in POLDIP3 and RANBP1 with TDP-43 knockdown and in TDPneg ALS/FTD neuronal nuclei. A. Venn diagram of genes with differential exon usage (DEU) common between HeLa, SH-SY5Y, ihMN and mouse striatum. B. Genes with exon usage changes in ALS/FTD TDP-43-immunonegative (TDPneg) neuronal nuclei that were shared with at least two experimental TDP-43 knockdown studies. C. DEXSeq plots of POLDIP3 from HeLa, SH-SY5Y, ihMN, mouse striatum and ALS/FTD neuronal nuclei. The boxed area indicates where splice changes are expected in *POLDIP3* with TDP-43 knockdown. D. IGV trace showing exon exclusion in *POLDIP3* exon 3 of the canonical transcript (purple boxed area) with TDP-43 knockdown in representative HeLa, SH-SY5Y and ihMN samples. Mouse striatum TDP-43 knockdown did not show equivalent exon usage change in *Poldip3*, but TDPneg ALS/FTD neuronal nuclei showed the same pattern of *POLDIP3* exon 3 skipping as the TDP-43 knockdown studies. E. DEXSeq plots of *RANBP1* from HeLa, SH-SY5Y and ALS/FTD neuronal nuclei. The boxed area indicates where splice changes are expected in RANBP1 with TDP-43 knockdown. F. IGV trace showing exon inclusion in *RANBP1* exon 3 of the canonical transcript (purple boxed area) with TDP-43 knockdown in representative HeLa, SH-SY5Y and ihMN samples. Mouse striatum with TDP-43 knockdown did not show equivalent exon usage change in *Ranbp1*, but TDPneg ALS/FTD neuronal nuclei showed the same pattern of *RANBP1* exon 3 inclusion as the TDP-43 knockdown studies.

In human cells, *TRAPPC12* exonic element E028 (within intron 7) was increased with TDP-43 knockdown, but not in TDPneg ALS/FTD neuronal nuclei, which instead showed increased usage of E047 (within exon 10) (Supplementary figure 1). The exon usage change in mouse *Trappc12* was not equivalent to either region (Fig. 5D). Indeed, mouse striatal TDP-43 knockdown samples showed exonic element usage (E005/ exon 9) that *decreased* compared to controls.

*POLDIP3* showed differentially used exons in all four TDP-43 knockdown studies and in TDPneg ALS/FTD neuronal nuclei. The skipped exon in TDP-43-knockdown HeLa, SH-SY5Y, and ihMN (Fig. 6C *upper*) is defined as exonic element E016 by *DEXSeq*, equivalent to ‘exon 3’ of the canonical transcript (ENSEMBL POLDIP3-201, ENST00000252115.10). *POLDIP3* exon 3 skipping following TDP-43 depletion was also shown previously (44). Although the mouse orthologue (*Poldip3*) also showed differential exon usage with TDP-43 knockdown, it was not in the equivalent region, instead occurring in exon 7 (Fig. 6C *middle*). Skipping of *POLDIP3* E016/ exon 3 was also observed in TDPneg ALS/FTD neuronal nuclei (Fig. 6C *lower*). In accordance with *DEXSeq* analysis, skipping of *POLDIP3* exon 3 was observed in all three TDP-43 knockdown human datasets (HeLa, SH-SY5Y, ihMN), and in TDPneg ALS/FTD neuronal nuclei, but not in TDP-43-knockdown mouse striatum, using IGV sashimi plots (Fig. 6D).

*RANBP1* showed alternative splicing of exon 3 in TDP-43-knockdown HeLa, SH-SY5Y, and ihMN (Fig. 6E). TDP-43-dependent splicing of this region was also reported previously, although it was annotated in those studies as exon 5 (47, 57). IGV sashimi plots again support both the positive and negative the findings from *DEXSeq* (Fig. 6F).

The neuronal gene *STMN2* harbors a cryptic exon that is usually suppressed by TDP-43 but is expressed following TDP-43 depletion (51, 52). Our re-analysis of that reported data from SH-SY5Y and ihMN cells confirmed the emergence of a cryptic exon between exons 1 and 2 in SH-SY5Y, ihMN, and TDPneg ALS/FTD neuronal nuclei (Fig. 7A). IGV sashimi plots (Fig. 7B) agreed with *DEXSeq*. They also demonstrated that *STMN2* cryptic exon expression was cell-type-specific as *STMN2* was not expressed in HeLa cells, and species-specific as the cryptic exon was not expressed at the equivalent position in mouse *Stmn2*.

**Figure 7.**
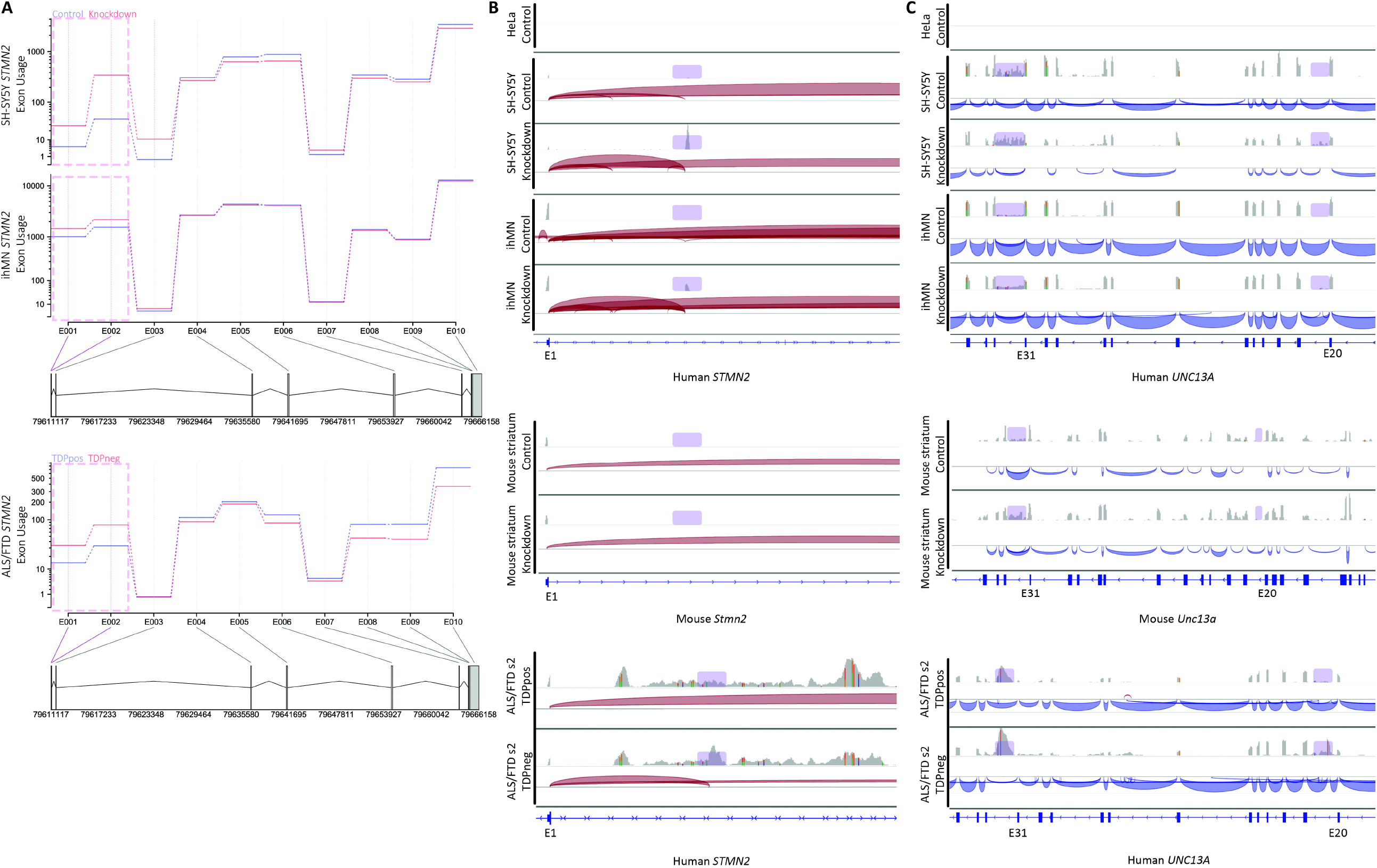
Known cryptic exons in STMN2 and UNC13A with TDP-43 knockdown and in TDPneg ALS/FTD neuronal nuclei. A. DEXSeq plots of STMN2 from SH-SY5Y, ihMN and ALS/FTD neuronal nuclei. The boxed area indicates where splice changes are expected in *STMN2* with TDP-43 knockdown. B. IGV trace showing cryptic exon inclusion in *STMN2* intron 1-2 of the canonical transcript (purple boxed area) with TDP-43 knockdown in representative SH-SY5Y and ihMN samples (*STMN2* not expressed in the HeLa cells). Mouse striatum with TDP-43 knockdown did not show cryptic exon expression in *Stmn2*, but TDPneg ALS/FTD neuronal nuclei showed the same cryptic exon as the TDP-43 knockdown studies. C. IGV trace showing *UNC13A* retained intron (intron 31-32, left purple box) and cryptic exon (in intron 20-21, right purple box) with TDP-43 knockdown in representative SH-SY5Y and ihMN samples. Mouse striatum IGV trace did not show the cryptic exon or retained intron but both were seen in the ALS/FTD neuronal nuclei IGV trace.

*UNC13A* is an ALS risk gene, in which expression of a cryptic exon between exons 20 and 21 and a retained intron between exon 31 and 32 was recently identified following TDP-43 knockdown (80, 81). Although neither region reached significance in our re-analysis using *DEXSeq*, when viewed with IGV (Fig. 7C) the cryptic exon and retained intron were enhanced with TDP-43 depletion in all human samples that expressed *UNC13A*, including TDPneg ALS/FTD neuronal nuclei. However, the cryptic exon was not observed in mouse striatum expressing *Unc13a* (Fig. 7C *middle*).

### Novel differentially used exons with TDP-43 knockdown and in TDPneg ALS/FTD neuronal nuclei

To our knowledge, TDP-43-dependent exon usage changes have not been previously highlighted in either *EXD3* or *CEP290. EXD3* showed cryptic exon expression in TDP-43-knockdown HeLa, SH-SY5Y, ihMN and TDPneg ALS/FTD neuronal nuclei (Fig. 8A). The cryptic exon is defined as exonic element E031 by *DEXSeq*, equivalent to ‘intron 7’ of the canonical transcript (ENSEMBL transcript EXD3-201, ENST00000340951.9). IGV sashimi plots clearly demonstrated read coverage of the intron in TDP-43-depleted samples (Fig. 8B).

**Figure 8.**
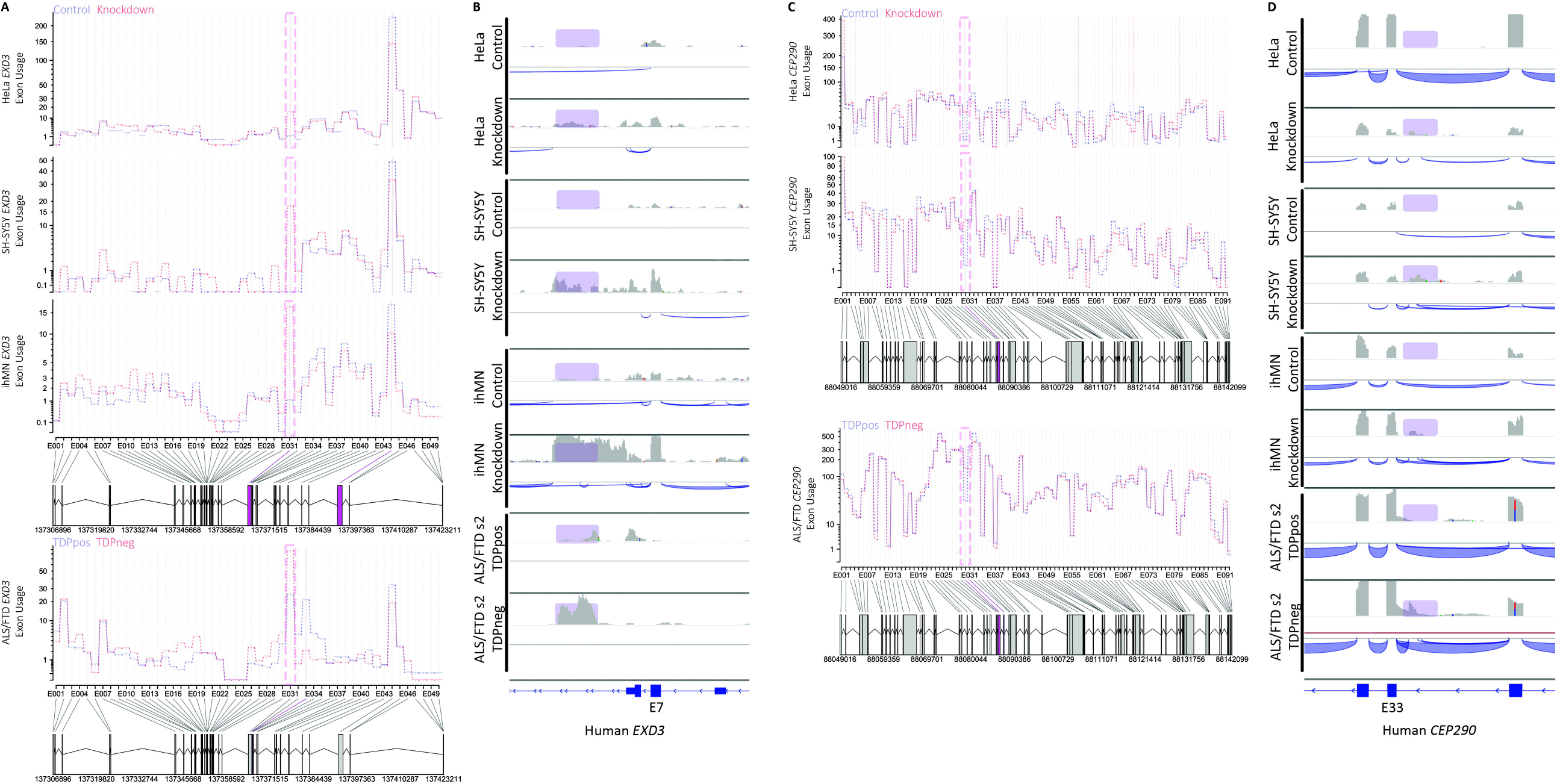
Novel cryptic exons in EXD3 and CEP290 with TDP-43 knockdown and in TDPneg ALS/FTD neuronal nuclei. A. DEXSeq plots of EXD3 from HeLa, SH-SY5Y, ihMN and ALS/FTD neuronal nuclei. The boxed area indicates where the cryptic exon occurs with TDP-43 knockdown. B. IGV trace showing cryptic exon inclusion in *EXD3* intron 7-8 of the canonical transcript (purple boxed area) with TDP-43 knockdown in representative HeLa, SH-SY5Y and ihMN samples. Representative TDPneg ALS/FTD neuronal nuclei show the same cryptic exon as the TDP-43 knockdown studies. C. DEXSeq plots of *CEP290* from HeLa, SH-SY5Y and ALS/FTD neuronal nuclei. The boxed area indicates where the cryptic exon occurs with TDP-43 knockdown. D. IGV trace showing cryptic exon inclusion in CEP290 intron 32-33 of the canonical transcript (purple boxed area) with TDP-43 knockdown in representative HeLa, SH-SY5Y and ihMN samples. A representative TDPneg ALS/FTD neuronal nuclei sample showed the same cryptic exon as the TDP-43 knockdown studies.

*CEP290* showed a cryptic exon in TDP-43-knockdown HeLa, SH-SY5Y, and TDPneg ALS/FTD neuronal nuclei (Fig. 8C). The cryptic exon is defined as exonic element E031 by *DEXSeq*, equivalent to ‘intron 32’ of the canonical transcript (ENSEMBL transcript CEP290-207, ENST00000552810.6). IGV sashimi plots supported the *DEXSeq* findings and also suggested cryptic exon emergence in the same intron in TDP-43-knockdown ihMN (Fig. 8D).

### Novel 3’ UTR usage with TDP-43 knockdown and in TDPneg ALS/FTD neuronal nuclei

Finally, we identified differential 3’ UTR usage in *KPNA4* and *MMAB*, which have not previously been described as TDP-43 target genes. *KPNA4* showed reduced 3’UTR coverage in TDP-43-knockdown HeLa, SH-SY5Y, ihMN and TDPneg ALS/FTD neuronal nuclei (Fig. 9A). The 3’ UTR of the transcript (ENSEMBL transcript KPNA4-001, ENST00000334256.4) is defined as exonic element E001 by *DEXSeq*. IGV sashimi plots showed a reduction in 3’ UTR reads in TDP-43-depleted samples (Fig. 9B).

**Figure 9.**
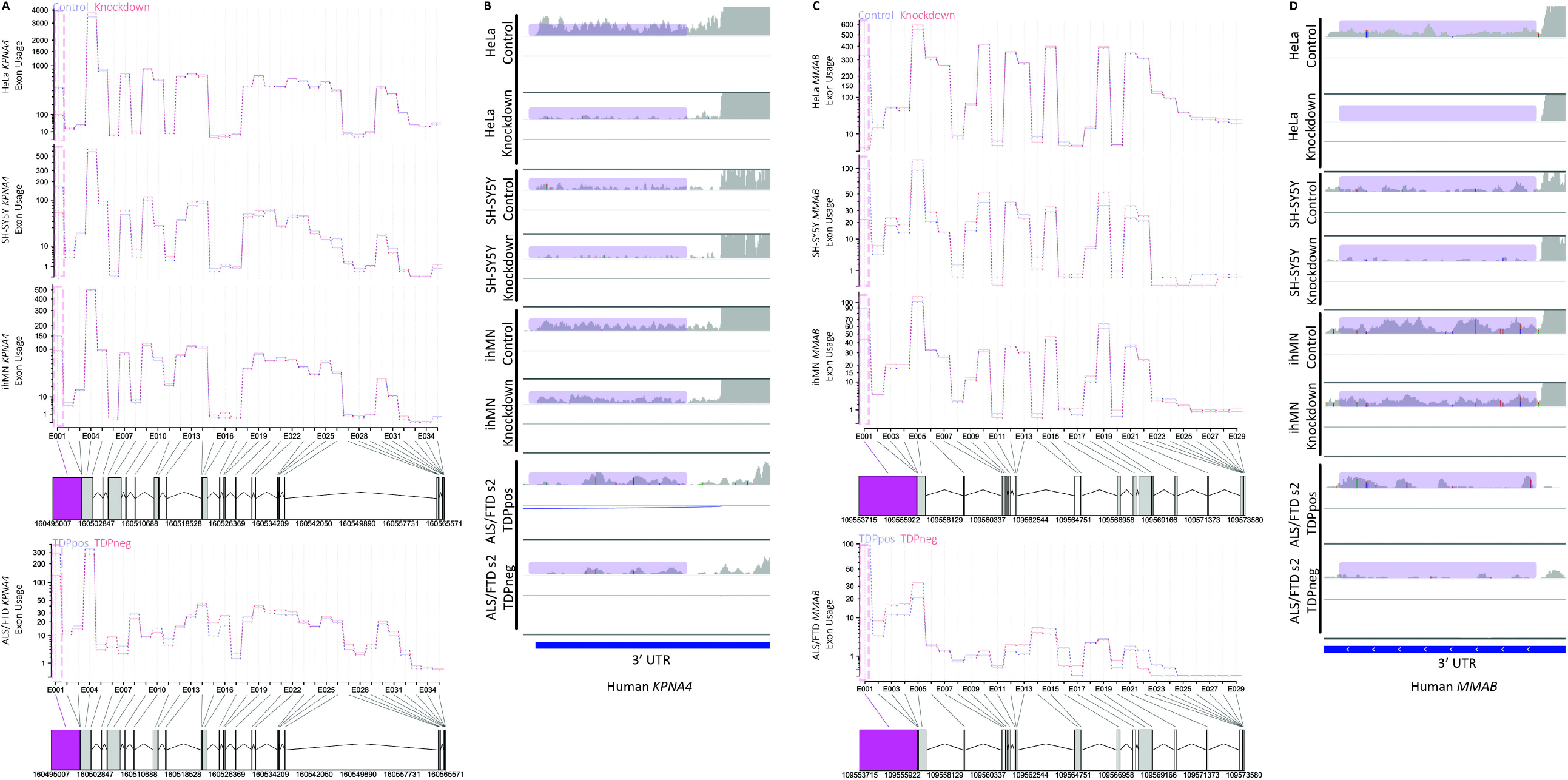
Novel differential 3’ UTR usage with TDP-43 knockdown and in TDPneg ALS/FTD neuronal nuclei. In addition to coding exons, TDP-43 knockdown also alters untranslated region (UTR) usage. A. DEXSeq plots of *KPNA4* from HeLa, SH-SY5Y, ihMN and ALS/FTD neuronal nuclei. The boxed area indicates the 3’ UTR of the gene. B. IGV trace showing altered 3’ UTR usage (purple boxed area) in *KPNA4* with TDP-43 knockdown in representative HeLa, SH-SY5Y and ihMN samples. Representative TDPneg ALS/FTD neuronal nuclei show the same changed 3’ UTR usage as the TDP-43 knockdown studies. C. DEXSeq plots of *MMAB* from HeLa, SH-SY5Y, ihMN and ALS/FTD neuronal nuclei. The boxed area indicates the 3’ UTR of the gene. D. IGV trace showing altered 3’ UTR usage (purple boxed area) in *MMAB* with TDP-43 knockdown in representative HeLa, SH-SY5Y and ihMN samples. A representative TDPneg ALS/FTD neuronal nuclei sample showed the same changed 3’ UTR usage as the TDP-43 knockdown studies.

Similarly, *MMAB 3’* UTR usage was suppressed in TDP-43-knockdown HeLa, SH-SY5Y, and TDPneg ALS/FTD neuronal nuclei (Fig. 9C). The 3’ UTR of the transcript (ENSEMBL transcript MMAB-210, ENST00000545712.7) is defined as exonic element E001 by *DEXSeq*. A striking reduction in *MMAB* 3’ UTR reads in TDP-43-depleted samples was evident using IGV sashimi plots (Fig. 9D).

## Discussion

As the common neuropathological feature in ALS with or without FTD, aggregated TDP-43 and its acquired toxic functions represent tractable molecular targets for therapy. Yet mounting evidence suggests that successful therapies will also require rescue of normal physiological functions of TDP-43 that are compromised in disease. Loss of gene expression and splicing regulatory function are now widely accepted features of ALS with TDP-43 proteinopathy, but how the specific complement of TDP-43 mRNA targets drives pathogenesis and phenotype remains unclear. Central to this, is the need to identify the complement of TDP-43 mRNA targets in distinct cell types and model systems, and to verify which of these targets is regulated by TDP-43 in human ALS/FTD neuronal nuclei. We approached this problem by assessing TDP-43 mRNA targets in human and mouse models, in neuronal and non-neuronal cell types, identifying shared and unique targets and biological pathways under TDP-43 control (Table 2). TDP-43 regulates certain common targets across myriad cell types, implying that TDP-43 proteinopathy in non-neuronal cells initiates processes that at least partially overlap with those occurring in neurons. Conversely, we identified cell-type-specific TDP-43 targets, indicating that the effects of loss of TDP-43 function are also partly unique to cell type. Notably, TDP-43-dependent changes identified in multiple model systems were validated to change in human ALS/FTD neuronal nuclei with TDP-43 nuclear depletion supporting TDP-43 loss of function as pathomechanistic.

### Cell-type independent TDP-43 targets

Certain transcripts were regulated by TDP-43 in myriad cell types, likely due to ubiquitous expression of TDP-43 and widespread expression of the target. The DEGs *PFKP, KIAA1324, HIGD1A, CHID1, RANBP1*, and *PKN1*, and the DEU targets *POLDIP3, RANBP1, EXD3*, and *CEP290*, all show low tissue specificity with transcripts expressed across at least 10 different organs (82) (Table 2*). S100A1* expression is group-enriched in brain and muscle, while KPNA4 is enriched in skeletal muscle and *MMAB* in liver (82).

Decreased *PFKP* mRNA has been identified as a TDP-43 loss-of-function marker by several studies in addition to those in our meta-analysis (56, 83, 84). *PFKP* encodes the platelet isoform of phosphofructokinase, a key glycolytic enzyme that is expressed almost ubiquitously. *PFKP* is proposed to decrease under conditions of TDP-43 depletion via suppression of miR-520 (84). However, TDP-43 can bind PFKP directly (47) and TARDBP knockdown results in *PFKP* cryptic exon exclusion supporting a direct interaction (51). Despite the apparent promise of *PFKP* as a TDP-43 loss-of-function marker, glycolysis is hypothesized to be a compensatory mechanism in ALS, and in human ALS spinal cord tissue with TDP-43 proteinopathy *PFKP* levels were found to increase rather than decrease compared to controls (85).

The skipping of exon 3 in *POLDIP3* is another frequently reproduced marker in TDP-43 knockdown studies (43, 44, 47, 57, 59, 86, 87), in which it undergoes an isoform switch. In the absence of TDP-43, the canonical variant 1 is decreased while variant 2 is increased due to exon 3 exclusion (43, 44). Interestingly, transfection of HEK293T cells with TDP-43 mutants p.Q331K and p.M337V also caused *POLDIP3* exon 3 exclusion, suggesting that mutant TDP-43 overexpression can induce loss of function of endogenous wildtype TDP-43 (88). *POLDIP3* protein (also known as S6K1 Aly/REF-like target (SKAR)) interacts with exon junctional complexes to increase the translation efficiency of spliced mRNAs (89). Our validation of *POLDIP3* exon 3 exclusion in human ALS/FTD neuronal nuclei with loss of nuclear TDP-43 strongly supports *POLDIP3* as a TDP-43 loss-of-function marker in ALS tissue, and indeed increased POLDIP3 variant 2 mRNA is seen in various motor regions of the CNS in ALS (44).

Our analyses also highlighted a number of TDP-43 targets that are less well studied in the context of ALS. The diverse cellular processes in which the protein products of these targets participate reflect, and may even underpin, some of the similarly diverse processes implicated in ALS (90, 91); nucleocytoplasmic transport (RANBP1 (92, 93), KPNA4 (94, 95), rotein degradation via autophagy (KIAA1324 (96)), mitochondrial homeostasis (HIGD1A (97)), and cytoskeletal regulation and axonal transport (PKN1 (98, 99), CHID1 (100), and S100a1 (101)).

The value of TDP-43-regulated transcripts that are expressed in many cell types as loss-of-function markers will depend upon the relative abundance of cells that develop TDP-43 loss of function, compared to cells that do not, in the tissue sampled. Furthermore, where TDP-43 loss-of-function causes a change in the abundance of a transcript but not its splicing, that transcript must be an upstream driver of change in functional pathways in ALS, and not be subject to secondary regulation as the pathway becomes dysfunctional, to be a useful loss-of-function marker. Arguably, specific splicing changes provide greater confidence than differential gene expression of TDP-43-dependent effects. To identify additional cell types with TDP-43 loss of function in ALS we therefore propose single cell analysis of splicing changes in genes with low tissue specificity *POLDIP3, RANBP1, EXD3*, and *CEP290*.

Although TDP-43 aggregates are found primarily in neurons and very rarely in non-neuronal cells such as glia or muscle (2, 102), TDP-43 function may yet be lost in cells without detectable aggregates through misfolding or nuclear clearing. The detection of TDP-43 loss-of-function markers in non-neuronal cells would support the notion that neurodegeneration in ALS is non-cell autonomous. Indeed, ALS-derived astrocyte models show changes in splicing (103), that could in part be due to TDP-43 dysfunction (104). The muscle-enriched transcript *KPNA4* encodes karyopherin-α3, which is widely reported to mediate nucleocytoplasmic trafficking deficits in *C9ORF72-*linked ALS (105-108). Suppression of *KPNA4* 3’ UTR usage represents i) an exciting mechanism by which TDP-43 loss of function might promote its own cytoplasmic mislocalisation in *C9ORF72*-linked ALS, and ii) a novel biomarker of TDP-43 loss of function in muscle.

### Neuronal TDP-43 targets

All neuronal models examined showed the signal transduction pathway to be regulated by TDP-43. Specific neuronal TDP-43 target transcripts were also identified in knockdown models that were validated in TDP-immunonegative ALS/FTD neuronal nuclei; *STMN2, ELAVL3*, and *UNC13A*. Loss of *STMN2* expression is associated with the emergence of a cryptic exon after exon 1, which may drive nonsense-mediated decay (109). This truncated *STMN2* mRNA was upregulated in the frontal cortex in FTD with TDP-43 pathology (109). STMN2 protein is essential for microtubule stability and thus cytoskeletal transport, synapse maintenance, and homeostasis of motor neurons. STMN2 has been identified as a key TDP-43 target in neurodegeneration, with deficits in axonal outgrowth and repair following TDP-43 depletion being rescued by restoration of STMN2 alone (51, 52). Cytoskeletal dynamics are further implicated in the pathogenesis of ALS/FTD by disease-causing mutations in *DCTN1* (110), *PFN1* (111, 112), and *TUBA4A* (113). Interestingly, ALS/FTD linked to any of these genes is associated with TDP-43 cytoplasmic aggregation and thus presumably TDP-43 nuclear clearing, raising the possibility of a feed-forward interaction between TDP-43 loss of function and cytoskeletal dysfunction. We predict the *STMN2* cryptic exon will be of high utility in detecting neuronal subtypes with TDP-43 loss-of-function. Further, expansion of a polymorphic intronic CA repeat in *STMN2* increases ALS risk and worsens disease (114), and a therapeutic that induces STMN2 for ALS is soon to enter clinical trial.

*ELAVL3* encodes a neuron-specific RNA-binding protein, and several studies in addition to those we analyzed indicate a depletion of ELAVL3 at both gene expression and protein level with TDP-43 knockdown (83, 86). *ELAVL3* downregulation has also been observed in ALS motor neurons (115, 116). The TDP-43-dependent cryptic exon reported to emerge in intron 3 (51, 115) was not recapitulated in another study (115) or in our re-analysis of TDP-43-negative ALS/FTD neuronal nuclei. As we note above, the most specific markers of TDP-43 loss-of-function are likely to be splicing changes, so further investigation of this cryptic exon in *ELAVL3* in ALS is warranted.

Lastly, *UNC13A* is a neuronal presynaptic protein involved in the fusion of neurotransmitter-containing vesicles with the plasma membrane (117, 118), and is critical to neurotransmission at the neuromuscular junction (119). A TDP-43-dependent retained intron in *UNC13A* (exon 31-32) was initially identified only in SH-SY5Y cells during our *DEXSeq* re-analysis, however inspection of reads using IGV revealed this retained intron also in ihMN and human TDPneg ALS/FTD neuronal nuclei. Read inspection also revealed the previously reported cryptic exon (exon 20-21) (80, 81) following TDP-43 knockdown and in TDP-43-negative ALS/FTD neuronal nuclei. The cryptic exon lies in intron 20, where two important *UNC13A* single nucleotide polymorphisms (SNPs) are situated; rs12608932 (C/C) is associated with increased susceptibility to ALS and increased risk of comorbid FTD with ALS (120-122), while rs12973192 is associated with increased ALS risk (123). Both *UNC13A* SNPs were found to increase susceptibility to exon 20-21 inclusion and retention of the distant intron 31 (80, 81). In our study, the cryptic exon was seen only in human and not at the equivalent location in mouse. Similarly, the retention of intron 31 was striking in human and ambiguous in mouse, in which there was high basal retention of the intron. A cryptic exon in mouse neuronal *Unc13a* has previously been reported following TDP-43 knockdown, that induces nonsense-mediated decay, however it lies between exons 1 and 2 (61). The discordance between human and mouse *UNC13A* regulation by TDP-43 may be critically important for preclinical drug testing, given that human variation in *UNC13A* predicts both shorter ALS survival and rescue of this shortened survival by lithium carbonate (121, 124, 125). Like *STMN2*, splicing of *UNC13A* holds great promise as a marker of TDP-43 loss of function in neurons.

### Methods and models for identifying TDP-43 targets

Several of our data suggest our *DESeq2* and *DEXSeq* analyses to be conservative methods for identifying DEGs and differential exon usage, and comparative studies of DEG packages have also demonstrated *DESeq2* to err on the conservative side (126, 127). For instance, exon usage changes in *UNC13A* that were evident in several datasets using IGV were not significant by *DEXSeq*. It is essential to identify consistent and reliable markers of TDP-43 loss of function to nominate targets with diagnostic or therapeutic potential. *DEXSeq* was chosen for this study for its conservative approach to controlling Type I error, leading to fewer false positives (69). TDP-43 may therefore regulate additional splicing events than those described here, and their identification could be aided by the combined use of multiple exon usage and splicing analysis tools. Conversely, exon usage changes identified in this study are unlikely to be due to sample variance and are indeed TDP-43-dependent.

In addition to the methodology used, the transcriptional targets of TDP-43 that we identified were dependent upon the species and fidelity of the cellular and animal models of ALS employed. While there was some overlap in TDP-43 loss-of-function mRNA and exon targets between human cell types, there was little overlap between human and mouse, even at the pathway level. The lack of common targets between mouse and human models may also be driven by the modest and variable mouse Tardbp knockdown in the mouse striatum by antisense oligonucleotides against TDP-43 (53), to avoid the lethality associated with embryonic *Tardbp* knockout (128, 129). In human cells, *TARDBP* depletion level predicted the degree of *UNC13A* cryptic exon inclusion, which in turn correlated with *STMN2* cryptic exon inclusion (80). Yet neither *Unc13a* nor *Stmn2* cryptic exons emerged with *Tardbp* depletion in mice. These results build upon the emerging consensus that TDP-43 has a distinct set of molecular targets in different cell types (61), suggesting many TDP-43 targets are species-specific. Human-derived transcriptomes are likely to be most suitable for identifying molecular pathways and drug targets relevant to human ALS.

The mechanism of modelling ALS is equally critical to identifying disease-relevant TDP-43 targets and pathways. Depleting TDP-43 is gaining acceptance in a field initially predominated by overexpression and TDP-43 mutant models (130, 131), and here we demonstrate that TDP-43 depletion recapitulates at least some of the transcriptional effects of loss of nuclear TDP-43 in ALS/FTD neuronal nuclei. TDP-43 knockdown can thus be considered an appropriate, even if partial, experimental paradigm of disease, for identifying mechanisms, biomarkers, and therapeutic targets.

## Conclusion

This study highlights transcripts whose expression or splicing can robustly report upon TDP-43 loss-of-function. We validate exon exclusion in *POLDIP3* and *RANBP1*, and cryptic exons in *EXD3* and *CEP290*, as markers of TDP-43 loss of function across a range of cell types. In contrast, 3’ UTR usage in *KPNA4* could report upon TDP-43 loss of function in muscle, and cryptic exon inclusion in *STMN2* and *UNC13A* are markers of TDP-43 loss of function in neurons. TDP-43 knockdown largely alters different biological pathways in human and mouse model systems, and human-derived models better recapitulate specific transcriptional and splicing changes that occur in ALS/FTD neuronal nuclei. Our findings enable the identification of non-neuronal cell types with TDP-43 loss of function, while revealing key players and pathways in the selective neuronal cell death that occurs in ALS and FTD.

## Table and figure legends

*Table 1.TDP-43 knockdown studies from NCBI GEO database*

*Table 2.Summary of genes and gene regions showcased as potential TDP-43 loss-of-function markers*

## Declaration

### Ethics approval and consent to participate

N/A

### Consent for publication

N/A.

### Availability of data and materials

All data generated or analyzed during this study are included in this published article and its supplementary information files.

### Competing interests

The authors declare that they have no competing interests.

### Funding

MC is supported by a University of Auckland Doctoral Scholarship. ELS is supported by Marsden FastStart and Rutherford Discovery Fellowship funding from the Royal Society of New Zealand [grant numbers 15-UOA-157, 15-UOA-003]. This work was also supported by grants from Motor Neuron Disease NZ, Freemasons Foundation of New Zealand, Matteo de Nora, Coker Family Trust, and PaR NZ Golfing. No funding body played any role in the design of the study, nor in collection, analysis, or interpretation of data nor in writing the manuscript.

### Authors’ contributions

Study design by MC, ELS. Data analysis and visualization by MC. Manuscript was written by MC and ELS. Both authors have read and approved the final manuscript.

## Acknowledgements

This publication is dedicated to the patients and families who contribute to our research. We also wish to acknowledge the use of New Zealand eScience Infrastructure (NeSI, https://www.nesi.org.nz) high performance computing facilities, funded jointly by NeSI’s collaborator institutions and the Ministry of Business, Innovation & Employment Research Infrastructure programme. We also thank Professor Mike Dragunow for helpful suggestions regarding the manuscript.

## Notes

https://mcao051.github.io/HeLa_Glimma/

https://mcao051.github.io/SH-SY5Y_Glimma/

https://mcao051.github.io/ihMN_Glimma/

https://mcao051.github.io/Mouse-striatum_Glimma/

https://mcao051.github.io/HeLa_DEU/

https://mcao051.github.io/SH-SY5Y_DEU/

https://mcao051.github.io/ihMN_DEU/

https://mcao051.github.io/Mouse_striatum_DEU/

https://mcao051.github.io/FTD_ALS_NeuN_DEU/

### Competing Interest Statement

The authors have declared no competing interest.

