## Supplementary material for "Novel and known transcriptional targets of ALS/FTD protein TDP-43: Meta-analysis and interactive graphical databases": Table

| **Genome** | **Cell/tissue type**  **sequenced** | **Method of TDP-43**  **knockdown** | **Library type** | **Accession code** | **Control vs. KD *n*** | **Ref** |
| --- | --- | --- | --- | --- | --- | --- |
| *Homo sapiens* | HUES3 differentiated into human motor neurons (ihMN) | Short interfering RNA (siRNA) | Paired | GSE121569 | 9 vs. 6 | (1) |
| *Homo sapiens* | SH-SY5Y | siRNA | Single | GSE122069 | 3 vs. 3 | (2) |
| *Homo sapiens* | HeLa-M | CRISPR-Cas9 | Paired | GSE136366 | 3 vs. 3 | (3) |
| *Mus musculus* | Mouse striatum | Antisense oligonucleotides | Single | GSE27218 | 11 vs. 10 | (4) |
| *Mus musculus* | Embryonic stem | *Tardbp* exon 3 floxed, removed upon Cre ­recombinase induction | Single | GSE21993 | 2 vs. 3 | (5) |
| *Mus musculus* | Mouse mammary gland | *Tardbp* exon 2 and 3 floxed mice bred with WAP-Cre transgenic mice | Paired | GSE116456 | 3 vs. 4 | (6) |
| *Rattus norvegicus* | Rat astrocytes | siRNA | Single | GSE135611 | 3 vs. 3 | (7) |
| *Homo sapiens* | NeuN-positive nuclei from FTD/ALS post-mortem tissue | NA (TDP-43-immunopositive versus TDP-43-immunonegative nuclei pools) | Paired | GSE126542 | 7 vs. 7 | (8) |

***Table 1. TDP-43 knockdown studies from NCBI GEO database***

|  | **Gene (and region)** | **Direction of change with TDP-43 depletion** | **Model that change occurs in**  (significant DEXSeq or observable IGV) | | | | | **Gene expression tissue specificity** |
| --- | --- | --- | --- | --- | --- | --- | --- | --- |
|  |  |  | **TDP-43 KD model** | | | | **TDPneg ALS/FTD**  **Neuronal nuclei** |  |
|  |  |  | **He** | **SH** | **MN** | **mSt** |  |  |
| **DEGs** | | | | | | | | |
|  | *PFKP* | Decreased |  |  |  |  |  | Non-specific |
|  | *ELAVL3* | Decreased |  |  |  |  |  | Neuronal |
|  | *STMN2* | Decreased |  |  |  |  |  | Neuronal |
|  | *KIAA1324* | Increased |  |  |  |  |  | Non-specific |
|  | *HIGD1A* | Increased |  |  |  |  |  | Non-specific |
|  | *CHID1* | Decreased |  |  |  |  |  | Non-specific |
|  | *RANBP1* | Decreased |  |  |  |  |  | Non-specific |
|  | *S100A1* | Increased |  |  |  |  |  | Non-specific |
|  | *PKN1* | Decreased |  |  |  |  |  | Non-specific |
| **DEU** | | | | | | | | |
|  | *POLDIP3* (exon 3) | Inclusion |  |  |  | * |  | Non-specific |
|  | *TRAPPC12* (intron 7) | Inclusion (CE) |  |  |  | * | * | Non-specific |
|  | *RANBP1* (exon 3) | Exclusion |  |  |  |  |  | Non-specific |
|  | *STMN2* (intron 1) | Inclusion (CE) |  |  |  |  |  | Neuronal |
|  | *UNC13A* (intron 20) | Inclusion (CE) |  |  |  |  |  | Neuronal |
|  | *EXD3* (intron 7) | Inclusion (CE) |  |  |  |  |  | Non-specific |
|  | *CEP290* (intron 32) | Inclusion (CE) |  |  |  |  |  | Non-specific |
|  | *KPNA4* (3’ UTR) | Exclusion |  |  |  |  |  | Non-specific |
|  | *MMAB* (3’UTR) | Exclusion |  |  |  |  |  | Non-specific |

***Table 2. Summary of genes and gene regions showcased as potential TDP-43 loss-of-function markers***

*change occurs in a different region of the gene

Abbreviations. DEGs, differentially expressed genes; DEU, differential exon usage; CE, cryptic exon; KD, knockdown; He, HeLa; SH, SH-SY5Y; MN, induced human motor neurons; mSt, mouse striatum
