## Supplementary figure 1 for "Novel and known transcriptional targets of ALS/FTD protein TDP-43: Meta-analysis and interactive graphical databases"

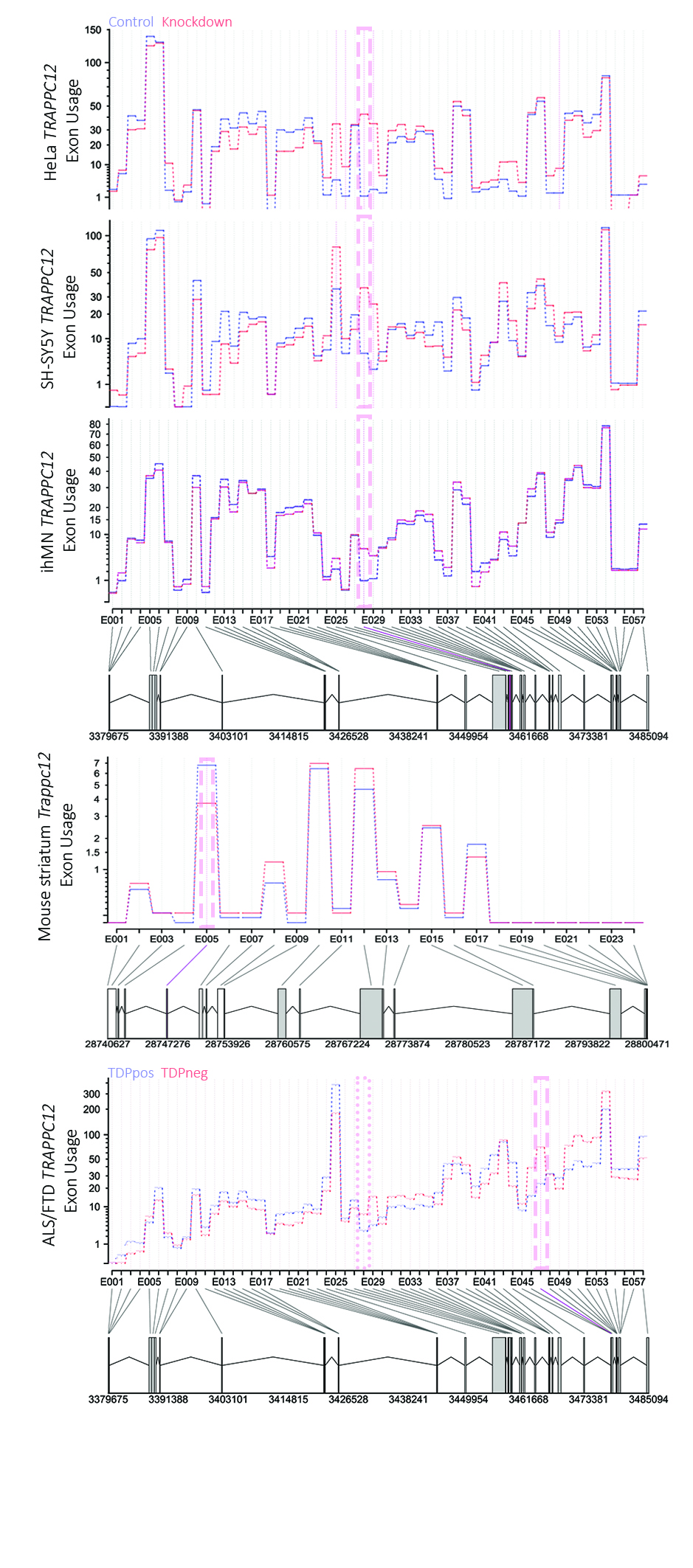

***Supplementary Figure 1.*** **DEXSeq exon usage plots for *TRAPPC12* in HeLa, SH-SY5Y, ihMN, mouse striatum and ALS/FTD neuronal nuclei.** Line-dashed boxes indicate significantly different exon usage for that dataset. The dot-dashed box in the ALS/FTD plot indicates no significant change in this dataset in the exonic region that was differentially used in human knockdown studies.
