## Supplementary file S1 for "Novel and known transcriptional targets of ALS/FTD protein TDP-43: Meta-analysis and interactive graphical databases"

#Code for trimming parameters

########################

\# GSE136366 (HeLa) example

########################

trimmomatic PE -threads $SLURM_CPUS_PER_TASK SRR10045018_1.fastq SRR10045018_2.fastq \

SRR10045018_1.trimmed.fastq SRR10045018_1un.trimmed.fastq \

SRR10045018_2.trimmed.fastq SRR10045018_2un.trimmed.fastq \

SLIDINGWINDOW:4:30 MINLEN:36

##########################

\# GSE122069 (SH-SY5Y) example

##########################

trimmomatic SE -threads $SLURM_CPUS_PER_TASK -phred33 \

SRR8144907.fastq \

SRR8144907.trimmed.fastq \

SLIDINGWINDOW:4:30 MINLEN:51

#########################

\# GSE121569 (ihMN) example

#########################

trimmomatic PE -threads $SLURM_CPUS_PER_TASK SRR8083864_1.fastq SRR8083864_2.fastq \

SRR8083864_1.trimmed.fastq SRR8083864_1un.trimmed.fastq \

SRR8083864_2.trimmed.fastq SRR8083864_2un.trimmed.fastq \

SLIDINGWINDOW:4:30 MINLEN:76

################################

\# GSE27218 (Mouse striatum) example

################################

trimmomatic SE -threads $SLURM_CPUS_PER_TASK -phred33 \

SRR107061.fastq \

SRR107061.trimmed.fastq \

ILLUMINACLIP:TruSeq2-SE.fa:2:30:10 SLIDINGWINDOW:4:30 MINLEN:72

###############################

\# GSE21993 (Mouse ES cells) example

###############################

trimmomatic SE -threads $SLURM_CPUS_PER_TASK -phred33 \

SRR052743.fastq \

SRR052743.trimmed.fastq \

ILLUMINACLIP:TruSeq2-SE:2:30:10 SLIDINGWINDOW:4:30 MINLEN:40

#######################################

\# GSE116456 (Mouse mammary gland) example

#######################################

trimmomatic PE -threads $SLURM_CPUS_PER_TASK SRR7455808_1.fastq SRR7455808_2.fastq \

SRR7455808_1.trimmed.fastq SRR7455808_1un.trimmed.fastq \

SRR7455808_2.trimmed.fastq SRR7455808_2un.trimmed.fastq \

SLIDINGWINDOW:4:30 MINLEN:150

##############################

\# GSE135611 (Rat astrocyte) example

##############################

trimmomatic SE -threads $SLURM_CPUS_PER_TASK -phred33 \

SRR9937050.fastq \

SRR9937050.trimmed.fastq \

ILLUMINACLIP:TruSeq3-SE.fa:2:30:10 SLIDINGWINDOW:4:30 MINLEN:151

#Code for *plotCount* graphs

###################

#View particular genes

###################

\# ‘dds’ is an object class from the *DESeq2* pipeline that stores read count information

which(rownames(dds)=="TARDBP")

tardbp <- plotCounts(dds, gene=rownames(dds)[52223], intgroup="condition",

returnData=TRUE)

p <- ggplot(tardbp, aes(x=condition, y=count)) +

ylim(0, NA) +

geom_boxplot(fill="orchid", colour="orchid4", width=0.5)

p + geom_dotplot(binaxis='y', stackdir='center', dotsize=0.5, fill="orchid4") +

ggtitle("TARDBP") +xlab("") + ylab("Normalized Count")+

theme_bw() +

theme(panel.grid.major = element_blank(), panel.grid.minor = element_blank(),text = element_text(size=20))

##################

\# View paired data

##################

library("ggbeeswarm")

which(rownames(dds)=="TARDBP")

tardbp <- plotCounts(dds, gene=rownames(dds)[52223], intgroup = c("condition","subject"),

returnData=TRUE)

ggplot(tardbp, aes(x = condition, y = count, color = subject, group = subject)) +

scale_y_log10() + geom_point(size = 3) + geom_line()
