## Supplementary file S2 for "Novel and known transcriptional targets of ALS/FTD protein TDP-43: Meta-analysis and interactive graphical databases"

**Interactive graphical databases of re-analyzed RNA-seq datasets**

*Glimma plots (differential gene expression):*

HeLa cells	<https://mcao051.github.io/HeLa_Glimma/>

SH-SY5Y cells	<https://mcao051.github.io/SH-SY5Y_Glimma/>

Induced human motor neurons	<https://mcao051.github.io/ihMN_Glimma/>

Mouse striatum	<https://mcao051.github.io/Mouse-striatum_Glimma/>

To **rank gene changes** by log_2_ fold-change (logFC) or adjusted p value (Adj.PValue), use arrows next to the parameter name in the gene table.

To **search for your gene of interest**, type the official gene symbol (GeneID) into the search box.

To **highlight a gene of interest on the scatter plot** comparing log_2_ fold-change with log mean expression, and generate a plot of expression of that gene for each sample (grouped by TDP-43 knockdown or nuclear clearing), double-click on the row for that gene in the gene table.

To **generate the plot of gene expression** for each sample/ group, hover over gene ‘dots’ in the scatter plot. To bring up the data for a gene in the gene table, double click the gene ‘dot’.

*DEXSeq plots (differential exon usage):*

HeLa cells	<https://mcao051.github.io/HeLa_DEU/>

SH-SY5Y cells	<https://mcao051.github.io/SH-SY5Y_DEU/>

Induced human motor neurons	<https://mcao051.github.io/ihMN_DEU/>

Mouse striatum	<https://mcao051.github.io/Mouse_striatum_DEU/>

NeuN-positive cells from ALS/FTD	<https://mcao051.github.io/FTD_ALS_NeuN_DEU/>

Please note that exonic elements, not exons, are examined and displayed graphically.

To **rank exonic element usage changes** by log_2_ fold-change (logFC) or adjusted p value (Adj.PValue), use Supplementary File S5.

To **search for the exonical element usage graphical display for your gene of interest**, use

Control+F (Windows) or Command+F (Mac) to find the entry for the Gene ID or ENSEMBL identifier and click the link.

The ‘counts’ tab gives normalized read counts per exonic element for each sample (best shows **variance**); the ‘expression’ tab gives mean exonic element expression for each sample group (best shows **DEGs**); ‘splicing’ tab gives mean exonic element usage for each sample group (best shows **differentially used exonic elements**); ‘transcripts’ tab gives mean exonic element expression for each sample group with respect to transcript variants (best shows **alternative splicing**); ‘results’ tab gives a table of raw data for each exonic element (best shows **raw data**).
